# Inspiratory muscle training in young, race-fit Thoroughbred racehorses during a period of detraining

**DOI:** 10.1101/843417

**Authors:** Lisa M. Katz, Jessica Stallard, Amy Holtby, Emmeline W. Hill, Kate Allen, James Sweeney

**Affiliations:** UCD School of Veterinary Medicine, University College Dublin, Belfield, Dublin 4, Ireland; Plusvital Ltd., Dun Laoghaire Industrial Estate, Pottery Road, Dublin, A96 KW29, Ireland; UCD School of Agriculture and Food Science, University College Dublin, Belfield, Dublin 4, Ireland; School of Veterinary Sciences, University of Bristol, Langford House, Langford, Bristol BS40 5DU, United Kingdom; University of Limerick, Ireland, Department of Mathematics & Statistics, Castletroy, Limerick, Ireland

## Abstract

Although inspiratory muscle training (IMT) is reported to improve inspiratory muscle strength in humans little has been reported for horses. We tested the hypothesis that IMT would maintain and/or improve inspiratory muscle strength variables measured in Thoroughbreds during detraining. Thoroughbreds from one training yard were placed into a control (Con, *n*=3 males *n*=7 females; median age 2.2±0.4 years) or treatment group (Tr, *n*=5 males, *n*=5 females; median age 2.1±0.3 years) as they entered a detraining period at the end of the racing/training season. The Tr group underwent eight weeks of IMT twice a day, five days per week using custom-made training masks with resistance valves and an incremental threshold of breath-loading protocol. An inspiratory muscle strength test to fatigue using an incremental threshold of breath-loading was performed in duplicate before (T_0_) and after four (T_1_) and eight weeks (T_2_) of IMT/no IMT using a custom-made testing mask and a commercial testing device. Inspiratory measurements included the total number of breaths achieved during the test, average load, peak power, peak volume, peak flow, energy and the mean peak inspiratory muscle strength index (IMSi). Data was analysed using a linear mixed effects model, *P*≤0.05 significant. There were no differences for inspiratory measurements between groups at T_0_. Compared to T_0_, the total number of breaths achieved (*P*=0.02), load (*P*=0.003) and IMSi (*P*=0.01) at T_2_ had decreased for the Con group while the total number of breaths achieved (*P*<0.001), load (*P*=0.03), volume (*P*=0.004), flow (*P*=0.006), energy (*P*=0.01) and IMSi (*P*=0.002) had increased for the Tr group. At T_2_ the total number of breaths achieved (*P*<0.0001), load (*P*<0.0001), volume (*P*=0.02), energy (*P*=0.03) and IMSi (*P*<0.0001) were greater for the Tr than Con group. In conclusion, our results support that IMT can maintain and/or increase aspects of inspiratory muscle strength for horses in a detraining programme.

## Introduction

Resistance training is a versatile form of exercise training with a large range of adaptations occurring including improved muscular strength, power, shortening velocity and endurance [1]. Respiratory muscle training (RMT) is a resistance training technique aimed towards improving the strength and function of the respiratory muscles using the general training principals of overload. It is claimed to be the most efficient way of improving respiratory muscle function [1].

RMT involves breathing against an increasing amount of resistance for a short period of time to overload the respiratory muscles, requiring them to work at a higher intensity and/or longer duration than normal [1]. RMT was first developed to assist people with breathing difficulties and alleviate symptoms of respiratory diseases such as chronic obstructive pulmonary disease (COPD) and asthma [2,3]. RMT has been reported to cause hypertrophy of ventilatory muscles with subsequent improvements in respiratory strength in both healthy people [4,5] and patients suffering from illness such as heart failure [6,7] and COPD [8,9].

The use of RMT in training and performance in humans has been extensively researched, with RMT theorised to strengthen respiratory muscles, making them more resistant to fatigue by slowing down and/or removing the negative influence of breathing on exercise tolerance [1]. RMT has been evaluated in several human sports including cycling [10–12], running [13] swimming [14] and rowing [15,16]. All studies found that only inspiratory muscle training (IMT) improves performance, with expiratory muscle training having little effect [1].

IMT has been found to benefit sprint performance in human athletes [1]. Human athletes who undergo a specific IMT programme before interval training have been reported to have an enhanced ability to train at a significantly harder level with greater improvements in their sprint performance when compared to a group who did not undergo IMT [17]. Furthermore, IMT has been shown to help sustain an athlete’s ability to repeatedly sprint [1]. Increased sprint speed and shortened active recovery breaks in-between sprint efforts have also been reported for athletes following IMT [18], with improvements identified after five to six weeks of IMT [19,20].

To date, there has only been one study investigating IMT reported in horses [21]. It was found that using customised equine face masks and commercially-available human IMT equipment, IMT and an incremental loading inspiratory muscle strength test (IMST) were well tolerated in a group of ten National Hunt racehorses, with higher values measured when horses were well acclimated to wearing the face masks [21]. The researchers reported that following eight weeks of IMT, the mean peak inspiratory muscle strength index (IMSi), an index representing the highest load at which a horse is able open the valve to complete a breath during the IMST, increased from 27 to 41 cmH_2_O [21].

The aim of the present study was thus to test the hypothesis that IMT could be used to maintain and/or improve measured inspiratory muscle strength variables in a group of race-fit Thoroughbred (Tb) racehorses during a period of detraining.

## Materials and Methods

### Sample population

The study was approved by University College Dublin Animal Research Ethics Committee with owner consent obtained for all procedures. Twenty Tb Flat racehorses from one training yard were selected for inclusion at the end of the 2017 racing season and/or training period. Horses were excluded if they had been out of exercise training under saddle for >6 weeks, were >4 years of age and/or were doing more than walking and/or trotting on an automated horse walker as exercise at the beginning of the study.

### Study groups

Horses were placed as equally as possible into either a treatment (Tr) or control (Con) group based on the date they finished exercise training, their exercise workload (box rest, walking, trotting [both using an automated horse walker]) at the time of entering the study, age and sex. Where a horse could not be fully sex- and age-matched between groups, the priority for distribution of horses between study groups was based on the amount of time out of exercise training and the exercise workload being undertaken when entering the study. The duration (days) and type of exercise (box rest, walking, trotting) was recorded for each horse for the duration of the study.

### Experimental protocol

#### Inspiratory muscle strength testing

Once placed into groups, all horses underwent a period of acclimatisation to the IMT and IMST equipment followed by an IMST performed before (T_0_) and after four (T_1_) and eight weeks (T_2_) of either IMT (Tr group) or no IMT (Con group). All IMT and IMST were performed by the same three people and occurred with each horse standing in a stable. Custom-made airtight masks covering the entire nose and held in place with a strap placed around the poll behind the ears were used for IMST (Fig 1) and IMT (Figs 2 and 3). Acclimatisation consisted of initially having an unfastened mask placed over the nose multiple times followed by the mask remaining fastened in place for two to four minutes; this was performed on multiple separate occasions over two to four days depending on each individual horse’s temperament. After at least two acclimatisation sessions with a mask alone, a training mask was fastened in place with a training valve set at the lowest resistance level inserted for a duration of four to eight breaths; this was to allow horses to become comfortable with the sensation of restricted breathing and the sound of air moving through the valve. Horses who reacted negatively during any of the acclimatisation sessions were removed from the study.

**Fig 1.**
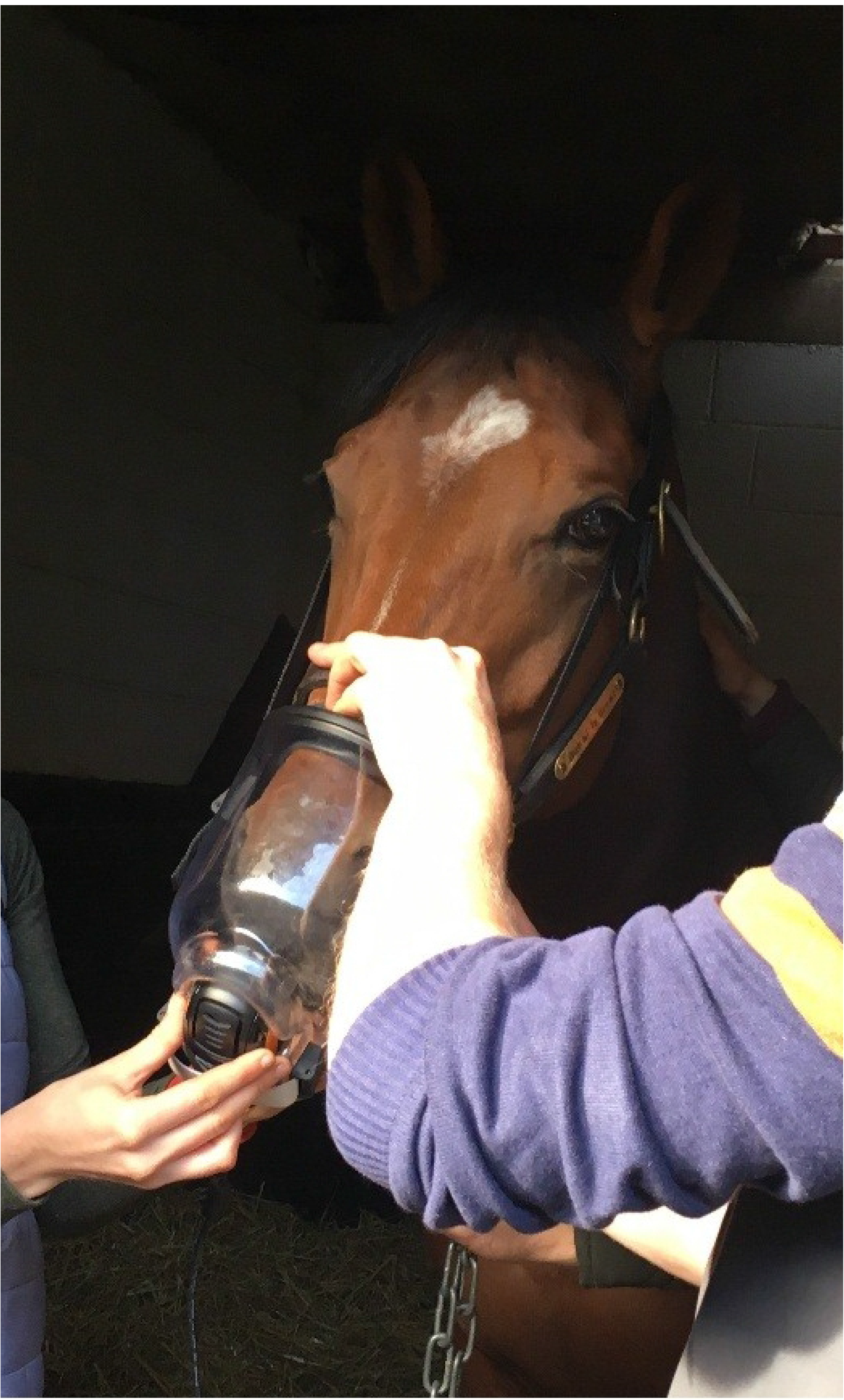
Inspiratory muscle strength testing to fatigue on a study subject.

**Fig 2.**
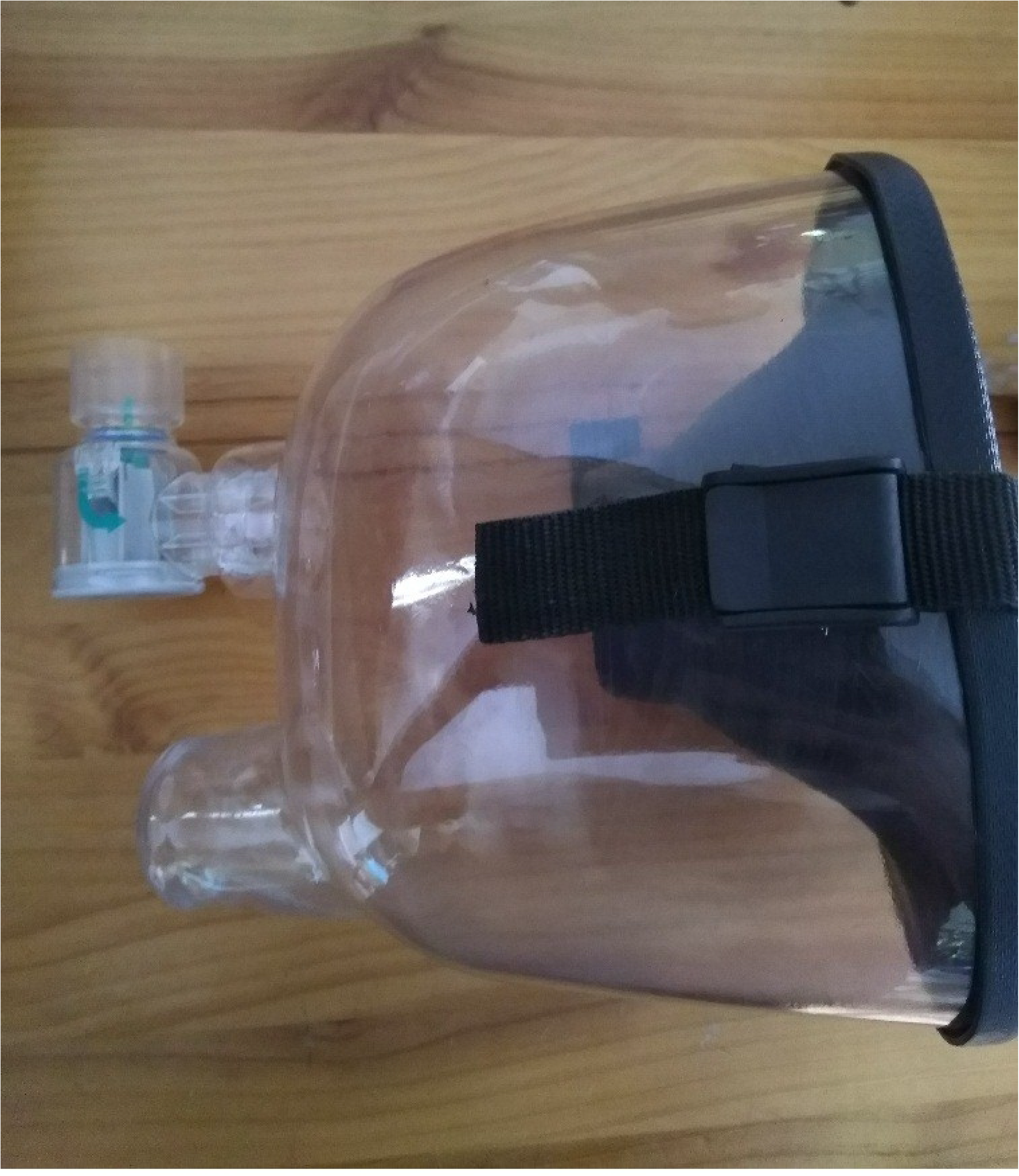
Training mask with a resistance valve inserted.

**Fig 3.**
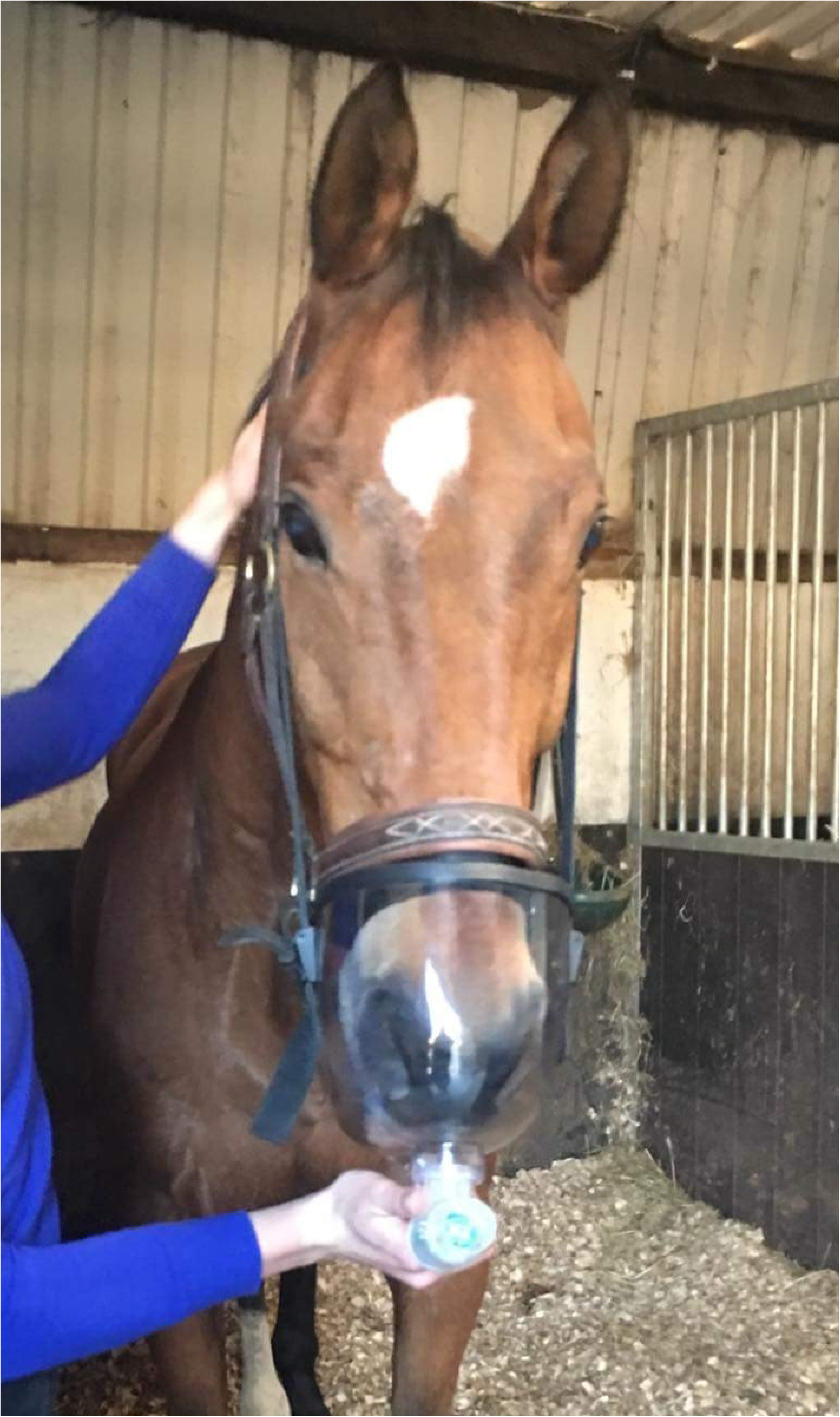
Training mask on a study subject with a resistance valve in place.

After the acclimatisation period, all horses underwent a baseline IMST in duplicate using a custom-made airtight testing mask and the POWERbreathe K5 (POWERbreathe International Ltd, UK), an electronic inspiratory loading device designed and validated for humans [22] and previously evaluated for use in horses [21]. The POWERbreathe K5 generates a pressure threshold which is required to be overcome for flow to occur. During the IMST, the total number of breaths achieved during the test, average load (i.e., inspiratory pressure, cmH_2_O), peak power (watts [W]), inspiratory volume (L), peak flow (L/sec) and energy (i.e., work of breathing, joules [J]) were continuously recorded at 500 Hz for each breath. The mean peak IMSi (cmH_2_O) was also recorded for each IMST. Customised software (Breathe-Link software, UK) was used for post-testing analysis of each testing session.

An incremental threshold of breath-loading protocol adapted from the human literature and developed for horses was used for the IMST [21] (Fig 4), in which the peak inspiratory pressure generated when breathing against an increasing amount of resistance is measured. The protocol consisted of an initial low threshold opening pressure, allowing the horse to become accustomed to the test, followed by an incremental loading protocol consisting of a ramp of increasing resistance to breathing by 2 cmH_2_O for each increment up to a potential maximum of 60 breaths in total. Each loaded breath was followed by two minimally loaded breaths (3 cmH_2_O), allowing the horse to recover before moving on to the next loaded breath. A loaded breath had to be completed to progress; if a horse failed twice to progress then the test ended.

**Fig 4.**
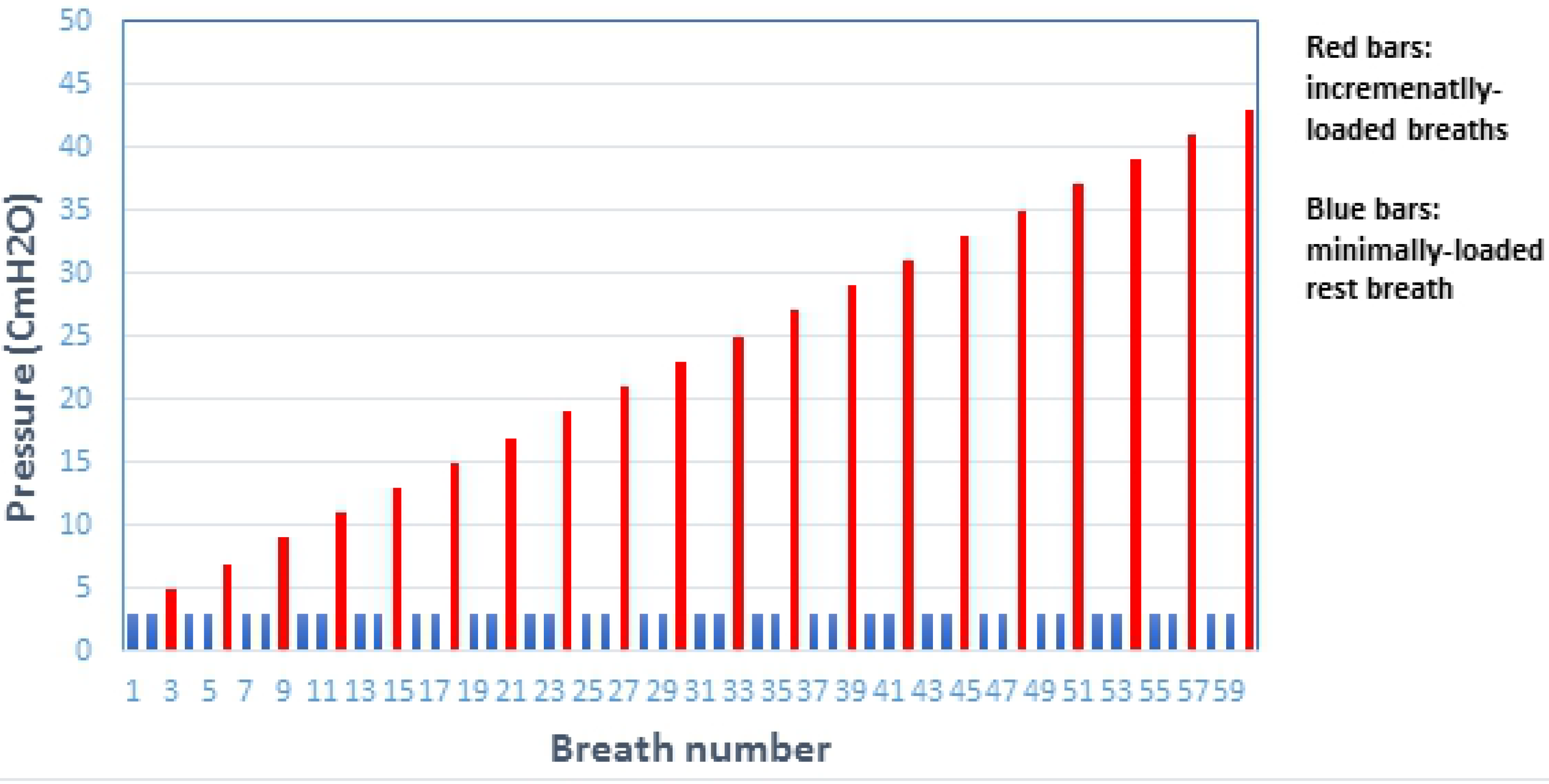
Incremental breath-loading protocol used for the respiratory muscle strength test to fatigue. Blue bars indicate low-resistance recovery breaths while red bars represent an increasing load of high-resistance breaths (modified from Allen et al., 2017).

#### Inspiratory muscle strength training

After baseline testing (T_0_), both groups of animals entered into an eight-week long study period with the Tr group receiving IMT twice a day, five days per week with a two to four-minute break between the two duplicate daily sessions. During this time, the Con group underwent weekly acclimatisation to the mask with a training valve as described previously to ensure similar familiarity and comfort with the mask as the Tr group during the study.

Each IMT session took between three to four minutes to complete, with the total daily IMT training (consisting of two duplicate IMT sessions) taking between 10—15 minutes each day to complete. Each IMT session consisted of 30 breaths against a pre-determined resistance to inhalation (i.e., loaded breaths) with the level of resistance at which a horse breathed against progressively increased over the training period. Resistance was created by using adjustable PEEP valves (Intersurgical, UK) that were inserted into one of two different custom-designed training masks (training mask 1 and training mask 2). Training mask 1 held wider diameter valves while training mask 2 fit to narrower diameter valves. The eight-week training protocol used had been previously established for use in horses [21], starting with a low resistance breathing load of 5 cmH_2_O followed by gradual increases in resistance every three to four days (10, 12.5, 15 and 20 cmH_2_O using training mask 1 followed by incremental increases of 2.5 cmH_2_O using training mask 2) until reaching a high-resistance breathing load of 40 cmH_2_O. The valves had been previously evaluated in conjunction with the training masks using a laboratory-based flow/volume simulator pump [Allen, unpublished] at flows and volumes reported for horses at rest [23,24], confirming that the valves created the prescribed inspiratory pressures and did not restrict airflow through flow resistance.

### Data analysis

All data analysis was conducted using the open source software package R [25]. All duplicate measures were included in the analysis. Values extracted for analysis included the overall peak value for a loaded or minimally loaded breath, and the peak value for the last loaded breath achieved during the IMST. The collected data for each variable of interest was treated as continuous with this assumption assessed via residual plots and the Shapiro-Wilk test. Where normality requirements were not satisfied due to skew tailed distributions, log transformations of the data were used. Categorical predictor data was assigned numerical values for analysis with exercise type during the study period assigned 0—2 (0=box rest, 1=walking, 2=trotting) and sex assigned 0 (male) or 1 (female). A linear mixed model with a random effect for each horse was used to assess the effect of IMT, duration of time out of exercise training (days) and the intensity (box rest, walking, trotting) and duration (days) of exercise during the study for each testing period (T_0_, T_1_, T_2_) on each measured respiratory variable (i.e., response variables). A linear relationship was assumed for the effect of each of the fixed effects on each response variable. A random effect for each horse was included in the model to account for the correlation in individual horse responses across both days and evaluations within days. Since each horse was evaluated in duplicate, evaluation order was assessed with no statistically significant difference identified in the response; evaluation order was thus excluded from the analysis.

Potential outliers for a given response were investigated by excluding them for an initial analysis and then examining the change in parameter estimates once they were included, using Student’s *t*-tests to assess for significant differences in the slope of each fixed effect. Multi-level categorical variables were assessed by ANOVA analysis. Results are expressed as the mean ± SEM with *P*≤0.05 significant.

## Results

The study groups are summarised in Table 1. Twenty-four horses were initially recruited with one horse dropped from the study due to inability to acclimatise to the mask and three additional horses dropped from the study due to being sold and leaving the training yard. The average days out of exercise training before entering the study were 14.9±12.8 days for the Tr group (*n*=5 males, *n*=5 females; median age 2.2±0.4 years) and 24.3±12.7 days for the Con group (*n*=3 males *n*=7 females; median age 2.2±0.4 years). All horses successfully completed the IMT programme, tolerating the testing and training masks well. The type of exercise for each group at each of the testing time-points are summarised in Table 2. The IMST results for each of the testing time-points are summarised in Table 3. Exercise and sex were not significantly associated with any of the measured inspiratory variables for either study group.

**Table 1.** Summary of a group of Thoroughbred horses (*n*=20) at the time of placement into an inspiratory muscle training treatment (*n*=10) or control (*n*=10) group based on when they finished exercise training and their current exercise workload.

| Treatment |  |  |  |  | Control |  |  |  |  |
| --- | --- | --- | --- | --- | --- | --- | --- | --- | --- |
| Horse | Sex | Age<br>(years) | Days out of<br>exercise training | Exercise workload<br>at study entry | Horse | Sex | Age<br>(years) | Days out of<br>exercise training | Exercise workload<br>at study entry |
| T01 | G | 2 | 27 | Walking/trotting | C01 | C | 2 | 15 | Walking/trotting |
| T02 | F | 2 | 27 | Walking | C02 | F | 2 | 25 | Walking |
| T03 | C | 2 | 24 | Box rest | C03 | F | 2 | 21 | Box rest |
| T04 | F | 2 | 27 | Walking | C04 | C | 2 | 42 | Walking |
| T05 | F | 2 | 11 | Walking/trotting | C05 | F | 2 | 15 | Walking/trotting |
| T06 | F | 2 | 5 | Walking | C06 | F | 2 | 28 | Walking |
| T07 | F | 3 | 0 | Walking/trotting | C07 | F | 3 | 5 | Walking/trotting |
| T08 | C | 2 | 0 | Trotting | C08 | C | 2 | 14 | Walking/trotting |
| T09 | C | 2 | 0 | Trotting | C09 | F | 2 | 36 | Walking/trotting |
| T10 | C | 2 | 28 | Walking/trotting | C10 | C | 3 | 42 | Walking/trotting |

Abbreviations: C: colt; G: gelding; F: female.

**Table 2.** The number of study horses at each exercise level in the inspiratory muscle training (IMT) treatment and control groups for each of the inspiratory muscle strength testing measurement time-points.

|  | Treatment (*n*=10) |  |  | Control (*n*=10) |  |  |
| --- | --- | --- | --- | --- | --- | --- |
| Time-points | box rest | walking<br>(on an automated<br>horse walker) | trotting<br>(on an automated<br>horse walker) | box rest | walking<br>(on an automated<br>horse walker) | trotting<br>(on an automated<br>horse walker) |
| T<sub>0</sub> | 2 | 6 | 2 | 2 | 4 | 4 |
| T<sub>1</sub> | 1 | 5 | 4 | 0 | 5 | 5 |
| T<sub>2</sub> | 1 | 3 | 6 | 0 | 4 | 6 |

Abbreviations: T<sub>0</sub>, before IMT; T<sub>1</sub>, after four weeks of IMT; T<sub>2</sub>, after eight weeks of IMT.

**Table 3.** The average peak measured values from the inspiratory muscle strength test to fatigue for each time-point for *n*=20 Thoroughbred horses in an inspiratory muscle training (IMT) treatment (Tr, *n*=10) or control (Con, *n*=10) group. Data presented as mean ± SEM and analysed using a mixed linear model with *P*≤0.05 significant. *significantly different from T_0_. ^§^significantly different from the Tr group.

| Time-points | Total breath number/test |  | IMSi (cmH<sub>2</sub>O) |  | Load (cmH<sub>2</sub>O) |  | Power (W) |  | Volume (L) |  | Flow (L/sec) |  | Energy (J) |  | Work of breathing (J/L) |  |
| --- | --- | --- | --- | --- | --- | --- | --- | --- | --- | --- | --- | --- | --- | --- | --- | --- |
|  | Tr | Con | Tr | Con | Tr | Con | Tr | Con | Tr | Con | Tr | Con | Tr | Con | Tr | Con |
| T<sub>0</sub> | 19.4<br>$\pm 1.4$ | 19.7<br>$\pm 1.9$ | 14.6<br>$\pm 0.9$ | 14.8<br>$\pm 1.3$ | 10.5<br>$\pm 0.7$ | 10.6<br>$\pm 0.8$ | 1.0<br>$\pm 0.1$ | 1.1<br>$\pm 0.2$ | 4.6<br>$\pm 0.4$ | 4.5<br>$\pm 0.5$ | 2.1<br>$\pm 0.2$ | 2.2<br>$\pm 0.3$ | 2.2<br>$\pm 0.2$ | 2.4<br>$\pm 0.5$ | 0.5<br>$\pm 0.03$ | 0.5<br>$\pm 0.1$ |
| T<sub>1</sub> | 18.4<br>$\pm 1.6$ | 15.6<br>$\pm 1.1$ | 13.9<br>$\pm 1.1$ | 12.1<br>$\pm 0.7$ | 9.4<br>$\pm 0.8$ | 8.7<br>$\pm 0.6$ | 0.9<br>$\pm 0.2$ | 1.0<br>$\pm 0.1$ | 5.0<br>$\pm 0.4$ | 4.8<br>$\pm 0.6$ | 2.1<br>$\pm 0.3$ | 2.0<br>$\pm 0.2$ | 2.4<br>$\pm 0.3$ | 2.4<br>$\pm 0.4$ | 0.5<br>$\pm 0.1$ | 0.5<br>$\pm 0.03$ |
| T<sub>2</sub> | 29.3<br>$\pm 2.8^*$ | 14.3<br>$\pm 1.0^{*\S}$ | 21.2<br>$\pm 1.9^*$ | 11.2<br>$\pm 0.7^{*\S}$ | 13.5<br>$\pm 0.9^*$ | 7.7<br>$\pm 0.5^{*\S}$ | 1.2<br>$\pm 0.1$ | 0.9<br>$\pm 0.2$ | 6.1<br>$\pm 0.2^*$ | 4.2<br>$\pm 0.7^{\S}$ | 2.8<br>$\pm 0.2^*$ | 2.1<br>$\pm 0.3$ | 3.4<br>$\pm 0.5^*$ | 2.0<br>$\pm 0.5^{\S}$ | 0.5<br>$\pm 0.1$ | 0.5<br>$\pm 0.02$ |

Abbreviations: T<sub>0</sub>, before IMT; T<sub>1</sub>, after four weeks of IMT; T<sub>2</sub>, after eight weeks of IMT; IMSi, inspiratory muscle strength index.

Intra-horse coefficient of variation (CV) was determined for IMST using one horse that underwent testing six times over one day (one test per hour, total of six hours): 15.6% for the number of breaths achieved, 13.3% for IMSi, 14.0% for load, 33.1% for power, 34.6% for volume, 29.6% for flow and 43.0% for energy. Inter-horse CVs were determined from all study horses at T_0_: 19.1% for the number of breaths achieved, 17.2% for IMSi, 15.7% for load, 27.6% for power, 31.8% for volume, 26.2% for flow and 39.9% for energy.

### Number of breaths achieved

The number of days since the last day of exercise training had a significant negative effect on the total number of breaths achieved during the IMST (*P*<0.001). There was no difference for the total number of breaths achieved between the Con and Tr groups at T_0_ (Table 3, Fig 5). However, there was a significant difference between the Con and Tr groups at T_2_ (*P*<0.0001), with the Tr group achieving a greater number of breaths than the Con group. Furthermore, the number of breaths achieved at T_2_ significantly decreased for the Con horses (*P*=0.02) and increased for the Tr horses (*P*<0.001) as compared to the T_0_ measurements (Table 3, Fig 5).

**Fig 5.**
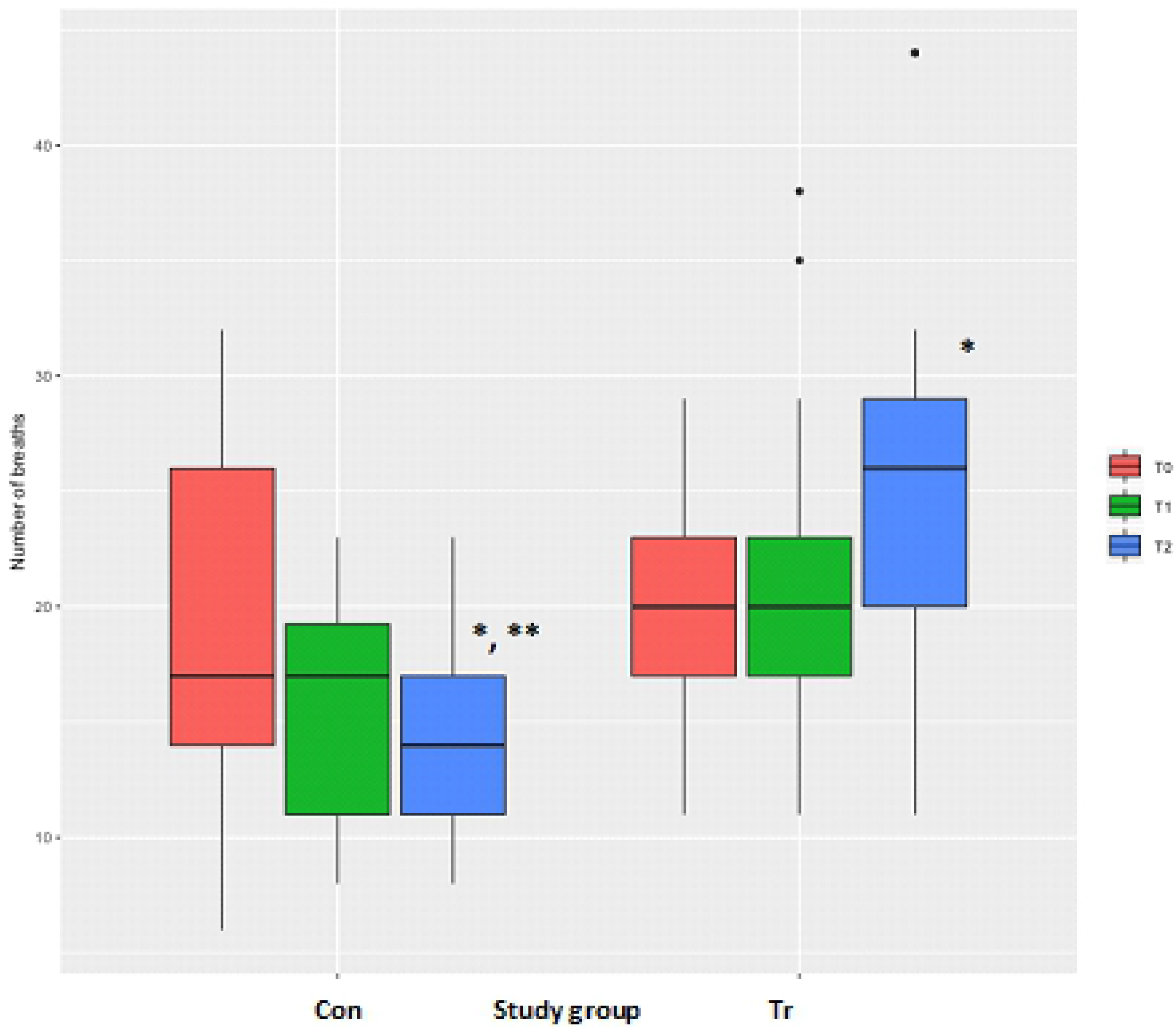
The average total number of inspiratory breaths achieved during inspiratory muscle strength testing to fatigue for each time-point for *n*=20 Thoroughbred horses in an inspiratory muscle training (IMT) treatment (Tr, *n*=10) or control (Con, *n*=10) group. Data presented as mean ± SEM and analysed using a mixed linear model with *P*≤0.05 significant. *significantly different from T_0_. **significantly different from the Tr group. T_0_: baseline, T_1_: after four weeks of IMT, T_2_: after eight weeks of IMT.

### Inspiratory muscle strength index

The number of days since the last day of exercise training had a significant negative effect on the peak IMSi achieved during the IMST (*P*=0.01). There were no differences for peak IMSi between the Con and Tr groups at T_0_ (Table 3, Fig 6). However, there was a significant difference between the Con and Tr groups at T_2_ (*P*<0.0001), with the Tr group able to breathe at a greater resistance than the Con group. At T_2_ the peak IMSi had significantly decreased for the Con group (*P*=0.01) and increased for the Tr group (*P*=0.002) as compared to T_0_ (Table 3, Fig 6).

**Fig 6.**
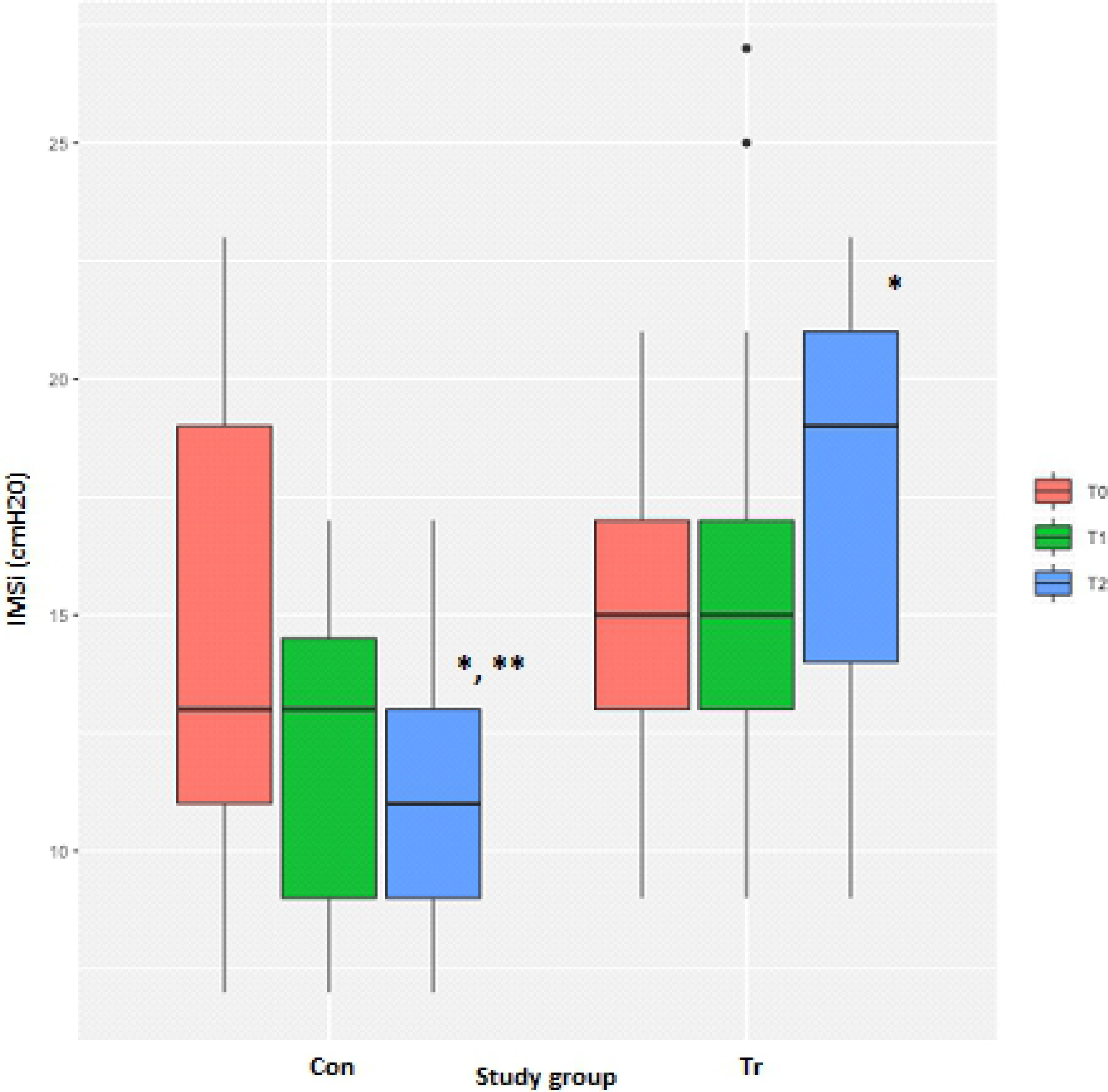
The average peak inspiratory muscle strength index (IMSi) measurements from inspiratory muscle strength testing to fatigue for each time-point for *n*=20 Thoroughbred horses in an inspiratory muscle training (IMT) treatment (Tr, *n*=10) or control (Con, *n*=10) group. Data presented as mean ± SEM and analysed using a mixed linear model with *P*≤0.05 significant. *indicates significantly different from T_0_. **indicates significantly different from the Tr group. T_0_: baseline, T_1_: after four weeks of IMT, T_2_: after eight weeks of IMT.

### Load (inspiratory pressure)

The number of days since the last day of exercise training had a significant negative effect while the duration of IMT training had a significant positive effect on the peak load achieved during the IMST (*P*=0.004). There were no differences for the peak load between the Con and Tr groups at T_0_ (Table 3, Fig 7). However, there was a significant difference for the peak load (*P*<0.0001) between the Con and Tr groups at T_2_, with the Tr horses able to achieve a greater load than the Con horses. The peak load that horses were able to achieve at T_2_ had significantly decreased in the Con group (*P*=0.003) and increased in the Tr group (*P*=0.03) as compared to T_0_ (Table 3, Fig 7).

**Fig 7.**
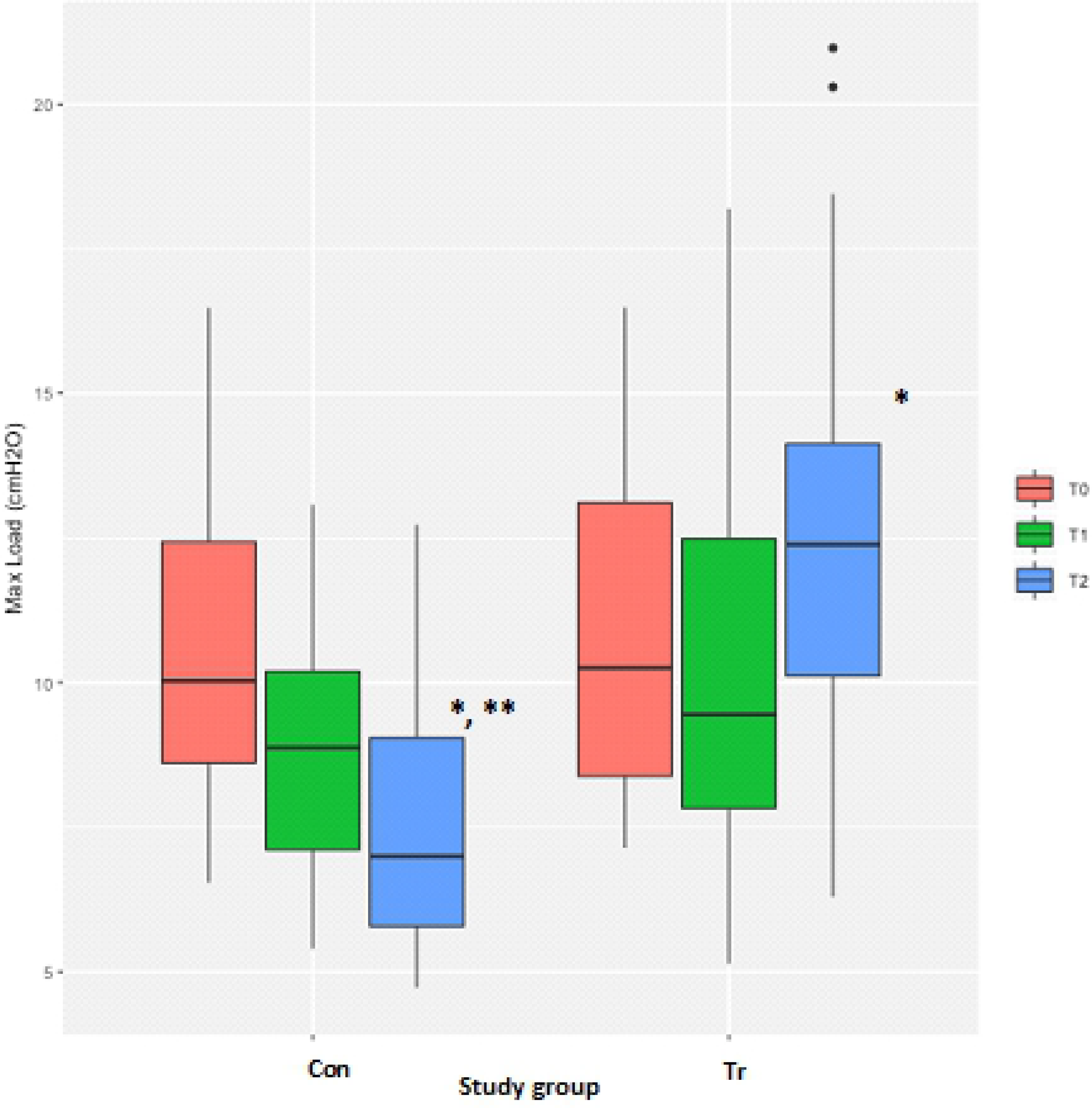
The average peak load measurements from inspiratory muscle strength testing to fatigue for each time-point for *n*=20 Thoroughbred horses in an inspiratory muscle training (IMT) treatment (Tr, *n*=10) or control (Con, *n*=10) group. Data presented as mean ± SEM and analysed using a mixed linear model with *P*≤0.05 significant. *indicates significantly different from T_0_. **indicates significantly different from the Tr group. T_0_: baseline, T_1_: after four weeks of IMT, T_2_: after eight weeks of IMT.

### Power

Measurements of the peak power achieved during the IMST were highly variable for both groups, with multiple extreme outliers identified. A log transformation of the power measurements successfully normalised the residuals after exclusion of observations with a power greater than 3.5 W. There were no differences for the peak power achieved during the test between the Con and Tr groups at T_0_ or at T_2_ (Table 3, Fig 8).

**Fig 8.**
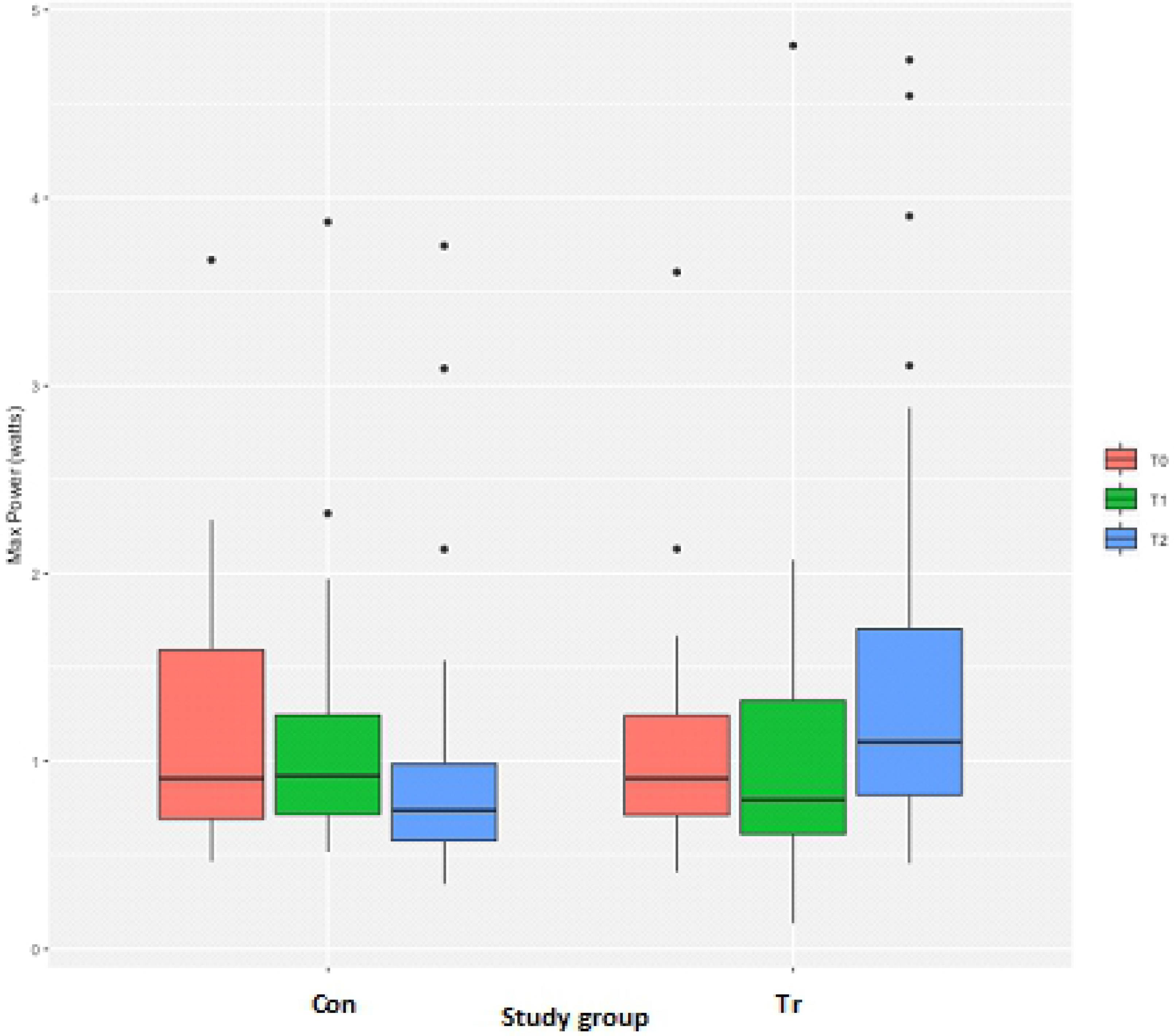
The average peak power measurements from inspiratory muscle strength testing to fatigue for each time-point for *n*=20 Thoroughbred horses in an inspiratory muscle training (IMT) treatment (Tr, *n*=10) or control (Con, *n*=10) group. Data presented as mean ± SEM and analysed using a mixed linear model with *P*≤0.05 significant. T_0_: baseline, T_1_: after four weeks of IMT, T_2_: after eight weeks of IMT.

### Volume

Measurements of the inspiratory volume achieved during the IMST were highly variable for both groups making it difficult to discern any patterns over time. A log transformation of the volume measurements successfully normalised the residuals. There were no differences for the peak volume achieved during the test between the Con and Tr groups at T_0_ (Table 3, Fig 9). However, there was a significant difference for the peak volume achieved between the Con and Tr groups at T_2_ (*P*=0.02), with the Tr horses able to achieve a greater volume during the test than the Con horses. There was no difference in the peak volume that horses in the Con group were able to achieve at T_2_ as compared to T_0_ (Table 3, Fig 9). Comparatively, the peak volume that horses in the Tr group were able to achieve at T_2_ had significantly increased as compared to T_0_ (*P*=0.004; Table 3, Fig 9).

**Fig 9.**
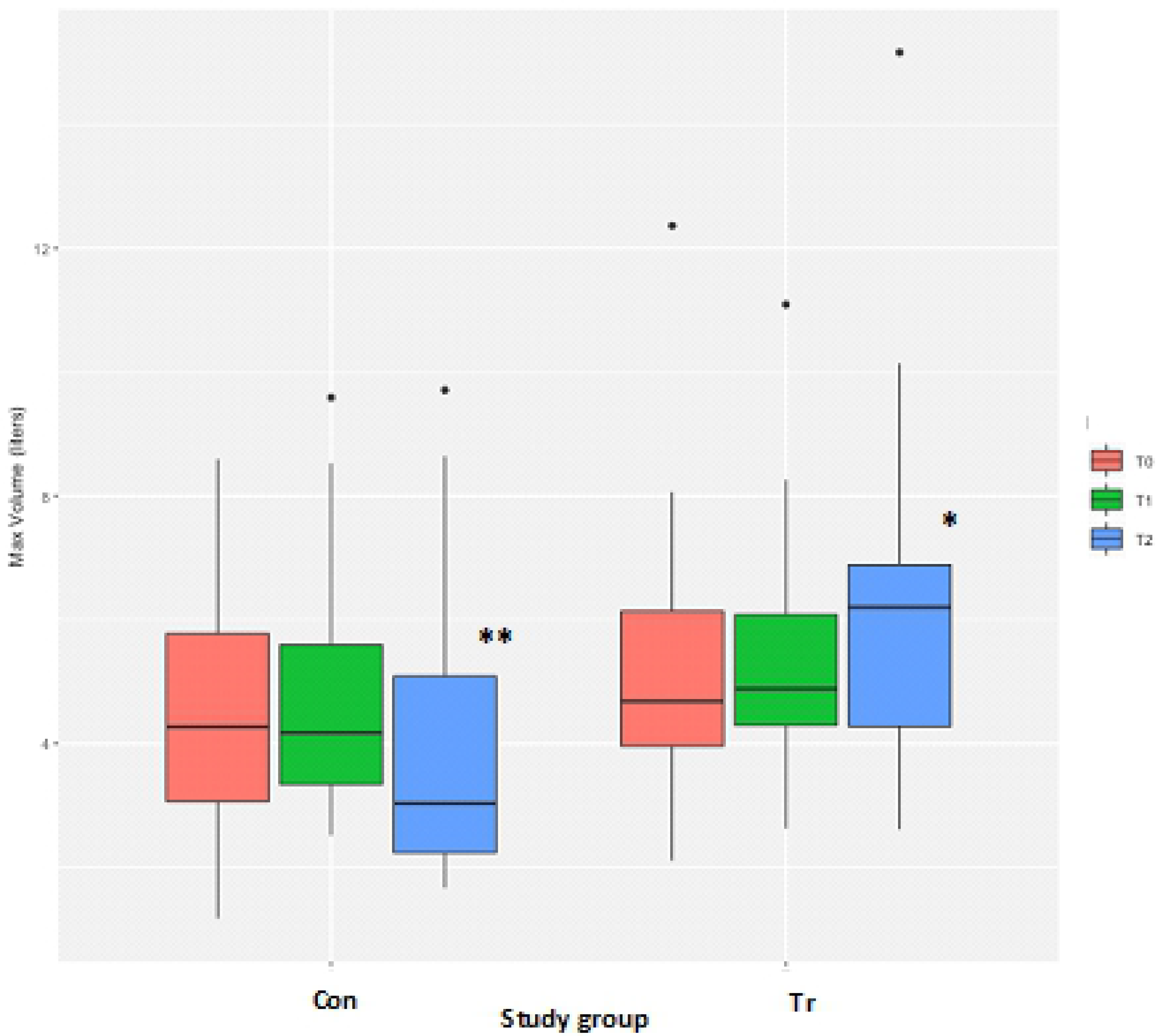
The average peak volume measurements from inspiratory muscle strength testing to fatigue for each time-point for *n*=20 Thoroughbred horses in an inspiratory muscle training (IMT) treatment (Tr, *n*=10) or control (Con, *n*=10) group. Data presented as mean ± SEM and analysed using a mixed linear model with *P*≤0.05 significant. *indicates significantly different from T_0_. **indicates significantly different from the Tr group. T_0_: baseline, T_1_: after four weeks of IMT, T_2_: after eight weeks of IMT.

### Flow

Measurements for peak flow achieved during the IMST were highly variable for both groups, although a log transformation of the data resulted in normalised residuals. There were no differences for the peak flow achieved between the Con and Tr groups at T_0_ (Table 3, Fig 10). There was no change in the peak flow that horses in the Con group were able to achieve at T_2_ as compared to T_0_ (Table 3, Fig 10). Comparatively, the peak flow achieved by Tr horses at T_2_ had significantly increased above T_0_ values (*P*=0.006; Table 3, Fig 10).

**Fig 10.**
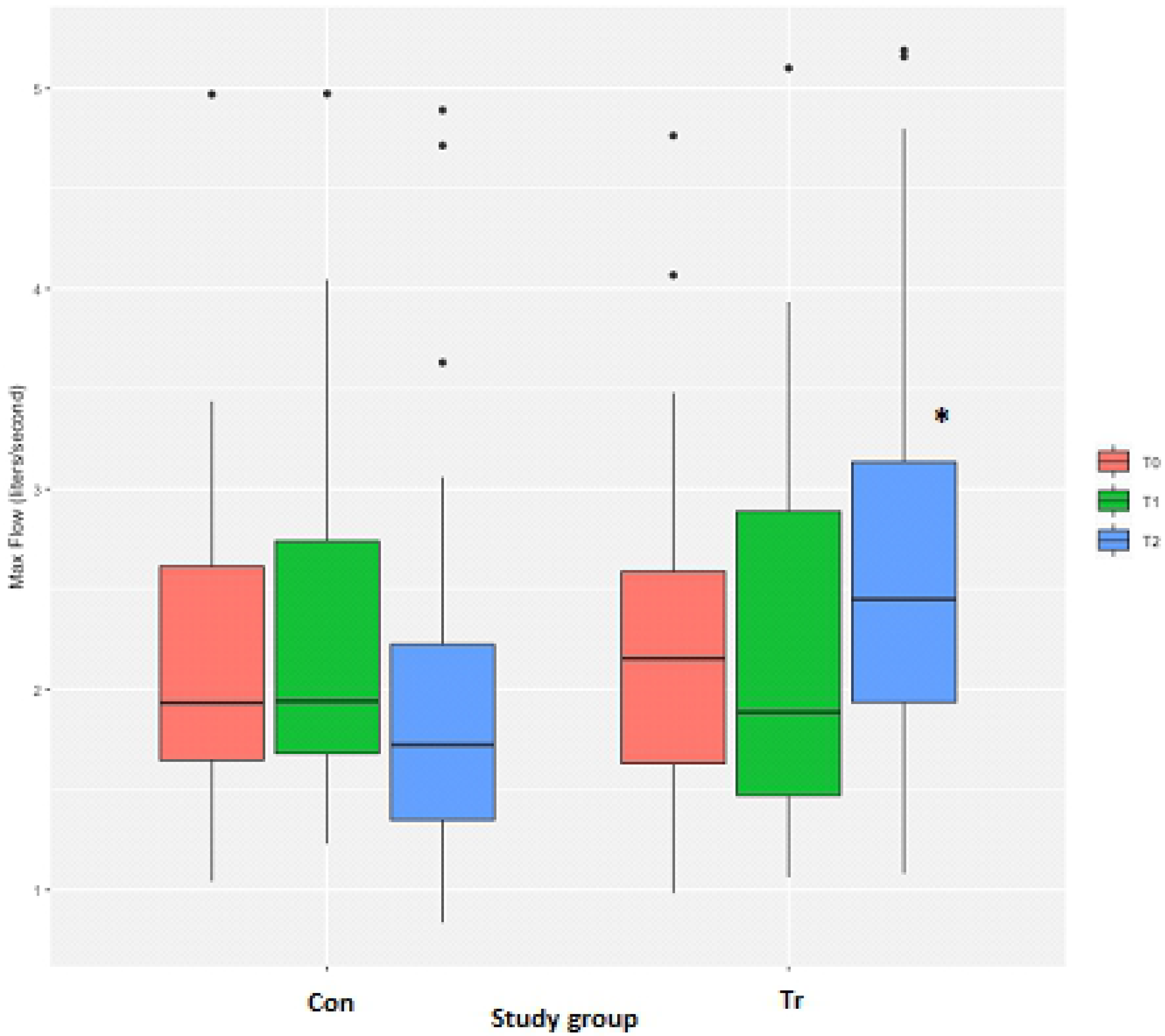
The average peak flow measurements from inspiratory muscle strength testing to fatigue for each time-point for *n*=20 Thoroughbred horses in an inspiratory muscle training (IMT) treatment (Tr, *n*=10) or control (Con, *n*=10) group. Data presented as mean ± SEM and analysed using a mixed linear model with *P*≤0.05 significant. *indicates significantly different from T_0_. T_0_: baseline, T_1_: after four weeks of IMT, T_2_: after eight weeks of IMT.

### Energy (work of breathing)

Measurements for the peak energy achieved during the IMST were highly variable, but a log transformation of the data and exclusion of observations greater than 7.5 J resulted in residuals closer to but not fully normalised, as the Shapiro-Wilk test still indicated non-normality of the residuals (*P*=0.01). There were no differences for the peak energy between the Con and Tr groups at T_0_ (Table 3, Fig 11). However, there was a significant difference between the Con and Tr groups at T_2_ (*P*=0.03), with the Tr group having higher energy measurements than the Con group. There was no change in the peak energy for the Con group at T_2_ as compared to T_0_ (Table 3, Fig 11). Comparatively, the peak energy for Tr horses at T_2_ had significantly increased above T_0_ values (*P*=0.01; Table 3, Fig 11).

**Fig 11.**
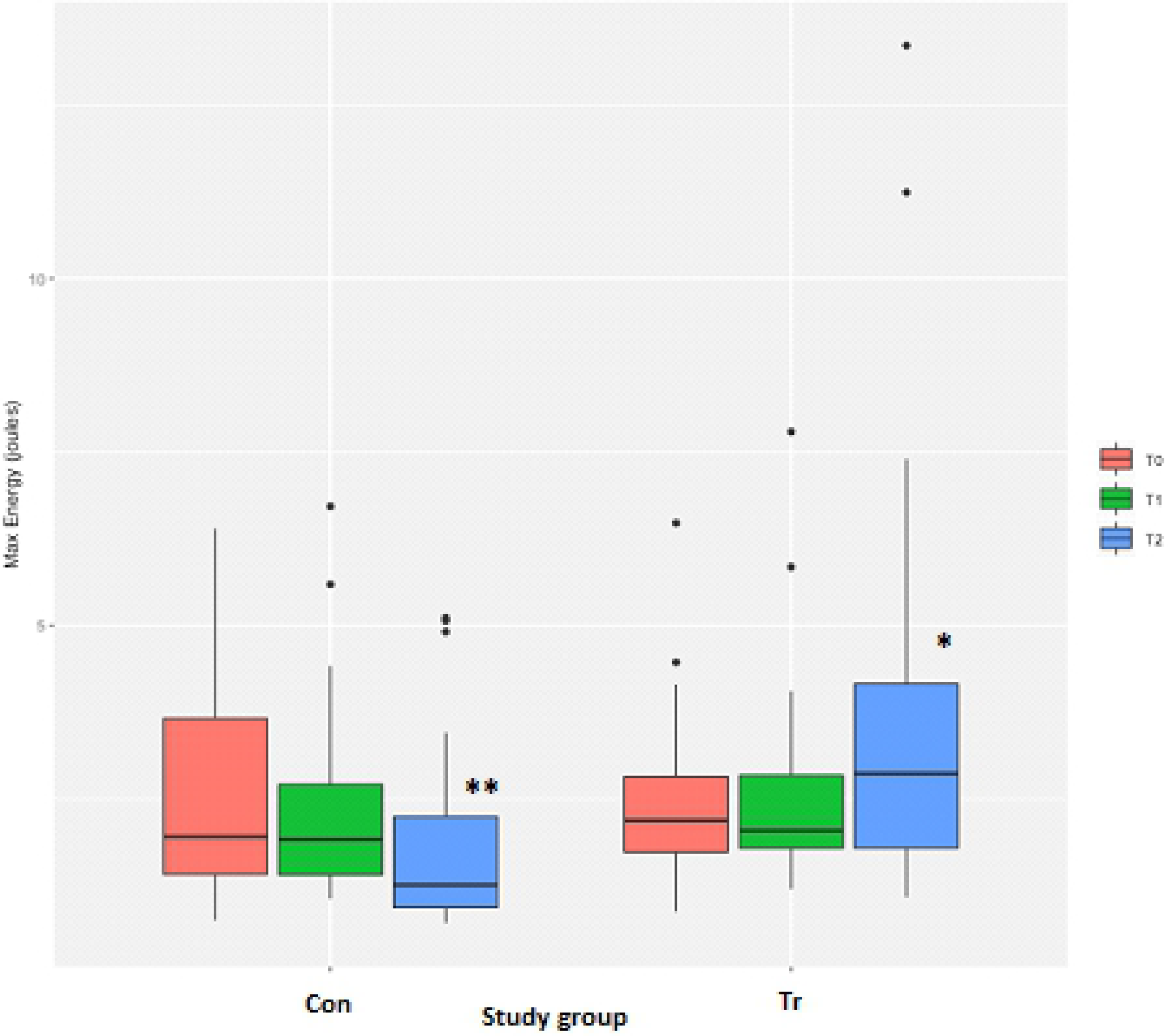
The average peak energy measurements from inspiratory muscle strength testing to fatigue for each time-point for *n*=20 Thoroughbred horses in an inspiratory muscle training (IMT) treatment (Tr, *n*=10) or control (Con, *n*=10) group. Data presented as mean ± SEM and analysed using a mixed linear model with *P*≤0.05 significant. *indicates significantly different from T_0_. **indicates significantly different from the Tr group. T_0_: baseline, T_1_: after four weeks of IMT, T_2_: after eight weeks of IMT.

## Discussion

This study is the first to demonstrate that a period of IMT affects resting inspiratory variables measured in race-fit Tbs during a period of detraining. Eight-weeks of IMT resulted in a significant increase above T_0_ values in the total number of breaths achieved, load, volume, flow, energy and ISMi variables measured during the IMST. Most of these variables measured after eight weeks of detraining (the total number of breaths achieved, load, volume, energy and IMSi) were also significantly greater for horses that had undergone IMT than those measured from horses that did not undergo IMT.

Horses were included in the study at the end of a racing/training season as they entered detraining, a period in which horses typically undertake minimal exercise activity as compared to their active exercise training programme. Human studies have shown that less fit athletes benefit more greatly from IMT than highly trained athletes [4,26,27], so the use of IMT as a training modality in horses during a period of relative inactivity was of interest. Furthermore, since privately owned horses were evaluated with no control over the training schedule, recruiting horses for the study as they entered a detraining period meant that all horses entered the study at relatively similar levels of fitness (race-fit) with similar levels of exercise (none/low-level) undertaken during the study. To further reduce the confounding effect of variation in fitness on the results, horses were primarily matched between study groups based on the duration of time out of exercise training and the type of exercise being undertaken when entering the study.

The type of IMT used in the present study was based on the principles of targeted resistance training of the inspiratory muscles, with horses standing in a stable and breathing against an increasing amount of flow-dependent resistance. Interestingly, ventilatory parameters measured during high-intensity exercise in horses have never been shown to improve in response to exercise training [28–30]. It has been hypothesised in humans that the respiratory muscles are not taxed in the same way the locomotory muscles are during exercise training, especially during short bouts of high-intensity training [31]. It is thus possible that this also occurs in horses, especially since the risk of musculoskeletal injury is quite high for horses exercising at high intensity with exercise training programmes having evolved to account for this risk. The typical Flat racehorse training programme uses a combination of lower-intensity exercise training with intermittent bouts of high-intensity sessions to provide a degree of anaerobic training while reducing the risk for musculoskeletal injury. As supported by the present results, IMT may be a way to allow the respiratory system to be overloaded with the horse at rest, inducing a measurable training response in the respiratory muscles.

After eight weeks of detraining, the total number of breaths, peak inspiratory load and IMSi (e.g., highest load at which a horse could open the test valve) achieved during the IMST significantly increased in the IMT group and decreased in the control horses. The total number of breaths and IMSi achieved during the IMST demonstrates the capability to inspire against an increasing load, reflected by the inspiratory pressure generated and associated force exerted by the respiratory muscles during inspiration. Since it is not possible to obtain maximal inspiratory efforts voluntarily from horses, a proxy for maximal inspiratory pressure (e.g., IMSi) was used as previously described [21]. Eight weeks of IMT also significantly increased the peak inspiratory volume, flow and energy measured during the IMST, with the latter reflecting the amount of mechanical work (e.g., effort) during inspiration. These results are all of interest since they may reflect increased strength and/or decreased metabolic demand of the inspiratory muscles during a certain level of work which could translate to decreased work of breathing during exercise.

In humans IMT has been demonstrated to increase the strength and endurance of the inspiratory muscles [1,17], possibly reducing the respiratory metabolic demands during exercise such that a greater proportion of available oxygen could be used by the locomotory muscles. This translates to enhanced athletic performance since increased work of breathing during exercise can reduce performance [32]. This is believed to be due to a greater percentage of the cardiac output being directed to the respiratory muscles as opposed to the locomotory muscles in order to meet metabolic demand [33], ultimately reducing energy production for locomotion [34]. In horses, the work of breathing is extremely high during high-intensity exercise due to the obligate nasal breathing and large minute ventilation generated to support the metabolic demand [23,35–38] with significant redistribution of blood flow from the skeletal locomotory muscles to the respiratory muscles [33,39]. It has also been shown that during high-intensity exercise the work of breathing in horses increases beyond a point (e.g., critical level of ventilation) [23,36,38,40] at which any additional oxygen made available by increases in ventilation is solely used for ventilation rather than locomotion. This is supported by the fact that the respiratory muscles of horses have extremely high oxidative capacities [41,42] and the ratio of the mechanical work of breathing to the amount of oxygen uptake (e.g., relative respiratory muscle oxygen uptake [23]) exponentially increases in conjunction with changes in minute ventilation [43].

It is also possible that IMT may improve performance in horses by reducing respiratory muscle fatigue as observed in humans [44–46]. Although respiratory muscles differ from other skeletal muscles because of their continuous activity, they have been shown to exhibit fatigue in humans following strenuous exercise [47,48], with diaphragmatic excitation-contraction decoupling occurring [49]. This has not yet been determined to occur in horses. Although there were no demonstratable effects of sex on the measurable outcomes in the present study, it is interesting to note that in humans the male diaphragm has been shown to be less fatigue resistant than the female diaphragm [50]. Respiratory muscle fatigue is primarily believed to negatively affect performance due to effects on the metaboreflex [48]. The metaboreflex refers to an accumulation of metabolites (i.e., lactic acid) within the respiratory muscles which have been linked with activation of group III and IV nerve afferents [51–53]. Activation of these nerve afferents have been shown to trigger increases in sympathetic outflow from the brain leading to vasoconstriction in the exercising limbs [54,55] and increased muscular limb fatigue [56,57].

Moderately high intra- and inter-horse variability for measurements obtained during the IMST support a degree of variation in how horses respond to IMT and IMST. Unlike IMT, the IMST is a test to fatigue with no way to ensure that each horse completes the test to fatigue versus just ‘giving up’, likely contributing to some of the observed variability. Since this was a pilot study with strict inclusion criteria used to standardise the study groups, the sample size was small as compared to IMT studies in humans. To fully evaluate the measured inspiratory variables and overcome the inherent intra- and inter-horse variability observed for the IMST, a larger sample population may be required.

It is relevant to note that highly variable baseline measures of inspiratory muscle strength between human individuals have been well documented [58,59]. None of the average baseline measurements in the present study significantly differed between the study groups but there was a degree of variation between individual horses within both study groups. The reason(s) behind these inter-subject variabilities are unknown but in humans, inherent differences between subjects in baseline inspiratory muscle strength [60] and the degree of activation of the diaphragm and chest wall inspiratory muscles during ventilation have been postulated [61]. Inherent differences in baseline strength of inspiratory muscles have also been shown to affect the efficacy of IMT in humans [62]. High inter-subject variability for the degree of improvement in maximal inspiratory pressure, an established measure of global inspiratory muscle strength in humans [63,64] following IMT has been demonstrated in numerous studies, with improvements ranging between 10—55% [15,44,65]. Although the IMSi (proxy for maximal inspiratory pressure) improved on average by 57% in horses following IMT, there was a wide degree of variation with improvements ranging between 29—146%. For the control group the IMSi decreased on average by 31% but there was also a wide degree of variation with decreases ranging between 11—57%. Since variability in limb skeletal muscle adaptation to strength training has been shown to be inversely related to baseline strength in humans [60], it is likely that the same applies for skeletal respiratory muscle adaptation to training. Although it was attempted to standardise fitness levels for all study horses, it is possible that a combination of variations in fitness as well as inherent differences in inspiratory muscle strength between horses contributed to the observed inter-horse variability in the present study.

Resting tidal volume values reported for horses are similar to T_0_ values obtained in the present study (4.8 L vs. 4.6 L), with reported resting peak inspiratory flows higher than the values measured in the present study (3.5 L/sec vs. 2.2 L/sec) [23,24]. Reported resting values for peak inspiratory pressure (0.2 cmH_2_O vs. 10.6 cmH_2_O), peak pulmonary resistance (0.21 cmH_2_O/L/sec vs. 14.7 cmH_2_O/L/sec) and peak work of breathing (0.41 J/L vs. 0.5 J/L) are all lower than the values measured in the present study [23,24]. This does make logical sense since values obtained for horses in the present study were obtained during an incremental threshold of breath-loading test which should result in greater inspiratory load, resistance and work of breathing values. However, since horses were used as their own controls, the purpose here was to determine the actual changes from baseline measurements rather than assess and interpret the actual measured values themselves.

## Conclusions

In conclusion, eight weeks of IMT significantly increased the total number of breaths achieved, load, volume, flow, energy and IMSi measured during an IMST in Tb Flat racehorses in a detraining programme. Comparatively, the total number of breaths achieved, load and IMSi significantly decreased below T_0_ values for control horses after eight weeks of detraining. These results support that IMT can be used to maintain and/or increase aspects of resting inspiratory muscle strength in horses not in active exercise training.

## Acknowledgements

The authors wish to express their appreciation to the trainer Mr. Jim Bolger and his staff, and in particular Mr Brian O’Connor and Mr Pat O’Donovan.

## References

1. McConnell A. Respiratory Muscle Training Theory and Practice. Great Britain: Churchill Livingstone; 2013.

2. Jones DT, Thomson RJ, Sears MR. Physical exercise and resistive breathing training in severe chronic airways obstruction-are they effective? Eur J Respir Dis. 1985;67(3):159–66.

3. McConnell A, Caine MP, Donovan KJ, Toogood AK, Miller MR. Inspiratory muscle training improves lung function and reduces exertional dyspnoea in mild/moderate asthmatics. Clin Sci. 1998;95:4P.

4. Illi SK, Held U, Frank I, Spengler CM. Effect of respiratory muscle training on exercise performance in healthy individuals: a systematic review and meta-analysis. Sports Med. 2012;42(8):707–24.

5. Hajghanbari B, Yamabayashi C, Buna T, Coelho JD, Freedman KD, Morton TA, et al. Effect of respiratory muscle training on performance in athletes: a systematic review with meta-analysis. J Strength Cond Res. 2013;6:1643–63.

6. Plentz RDM, Sbruzzi G, Ribeiro RA, Ferreira JB, Dallago P. Inspiratory muscle training in patients with heart failure: meta-analysis of randomized trials. Arq Bras Cardiol. 2012;99(2):762–71.

7. Smart NA, Giallauria F, Dieberg G. Efficacy of inspiratory muscle training in chronic heart failure patients: a systematic review and meta-analysis. Int J Cardiol. 2013;167(4):1502.

8. Geddes EL, O’Brien K, Reid WD, Brooks D, Crowe J. Inspiratory muscle training in adults with chronic obstructive pulmonary disease: An update of a systematic review. Respir Med. 2008;102(12):1715–29.

9. Gosselink R, De Vos J, Van Den Heuvel SP, Segers J, Decramer M, Kwakkel G. Impact of inspiratory muscle training in patients with COPD: what is the evidence? Eur Respir J. 2011;37(2):416–25.

10. Caine MP, McConnell AK. Pressure threshold inspiratory muscle training improves submaximal cycling performance. Third Annual Conference of the European College of Sport Science. Centre for Healthcare Development, Manchester, 1998.

11. Gething AD, Williams M, Davies B. Inspiratory resistive loading improves cycling capacity: a placebo-controlled trial. British J Sports Med. 2004;38(6):730–6.

12. Bailey SJ, Romer LM, Kelly J, Wilkerson DP, Dimenna FJ, Jones AM. Inspiratory muscle training enhances pulmonary O(2) uptake kinetics and high-intensity exercise tolerance in humans. J Appl Physiol. 2010;109(2):457–68.

13. Edwards AM, Cooke CB. Oxygen uptake kinetics and maximal aerobic power are unaffected by inspiratory muscle training in healthy subjects where time to exhaustion is extended. Eur J Appl Physiol. 2004;93(1-2):139–44.

14. Kilding AE, Brown S, McConnell AK. Inspiratory muscle training improves 100 and 200 m swimming performance. Eur J Appl Physiol. 2010;108(3):505–11.

15. Vollianitis S, McConnell AK, Koutedakis Y, McNaughton L, Backx K, Jones DA. Inspiratory muscle training improves rowing performance. Med Sci Sports Exerc. 2001;33(5):803–9.

16. Griffiths LA, McConnell AK. The influence of inspiratory and expiratory muscle training upon rowing performance. Eur J Appl Physiol. 2007;99(5):457–66.

17. Tong TK, Fu FH, Eston R, Chung P-K, Quach B, Lu K. Chronic and Acute Inspiratory Muscle Loading Augment the Effect of a 6-Week Interval Program on Tolerance of High-Intensity Intermittent Bouts of Running. J Strength Cond Res. 2010;24(11):3041–48.

18. Bangsbo J, Iaia FM, Krustrup P. The Yo-Yo intermittent recovery test: a useful tool for evaluation of physical performance in intermittent sports. Sports Med. 2008;38(1):37–51.

19. Tong TK, Fu FH. Effect of specific inspiratory muscle warm-up on intense intermittent run to exhaustion. Eur J Appl Physiol. 2006;97(6):673–80.

20. Nicks CR, Morgan DW, Fuller DK, Caputo JL. The influence of respiratory muscle training upon intermittent exercise performance. Int J Sports Med. 2009;30(1):16–21.

21. Allen K, McConnell A, Franklin S, Fitzharris L. Development of an inspiratory muscle testing and traingin device in the horse: a feasability trial. WEAS 2017.

22. Langer D, Jacombe C, Charususin N, Scheers H, McConnell A, Decramer M, et al. Measurement validity of an electronic inspiratory loading device during a loaded breathing task in patients with COPD. Respir. Med. 2013;107:633–635.

23. Art T, Anderson L, Woakes AJ, Roberts C, Butler PJ, Snow DH, et al. Mechanics of breathing during strenuous exercise in Thoroughbred horses. Respir Physiol. 1990;82(3):279–94.

24. Butler PJ, Woakes AJ, Smale K, Roberts CA, Hillidge CJ, Snow DH, et al. Respiratory and cardiovascular adjustments during exercise of increasing intensity and during recovery in thoroughbred racehorses. J Ex Biol. 1993;179:159–80.

25. R Core Team. R: a language and environment for statistical computing. R foundation for statistical computing, Vienna, Austria; 2016. Available at http://www.R-project.org/, 1–12.

26. Coast JR, Clifford PS, Henrich TW, Stray-Gundersen J, Johnson RL. Maximal inspiratory pressure following maximal exercise in trained and untrained subjects. Med Sci Sports Exerc. 1990;22(6):811–5.

27. Choukroun ML, Kays C, Gioux M, Techoueyres P, Guenard H. Respiratory muscle function in trained and untrained adolescents during short-term high intensity exercise. Eur J Appl Physiol Occup Physiol. 1993;67(1):14–9.

28. Art T, Lekeux P. Training-induced modifications in cardiorespiratory and ventilatory measurements in Thoroughbred horses. Equine Vet J. 1993;25(6):532–6.

29. Christley RM, Hodgson DR, Evans DL, Rose RJ. Effects of training on the development of exercise-induced arterial hypoxemia in horses. Am J Vet Res. 1997;58(6):653–57.

30. Roberts, CA, Marlin DJ, Lekeux P. The effects of training on ventilation and blood gases in exercising Thoroughbreds. Equine Vet J Suppl. 1999;31:57–61.

31. McConnell A. Breathe Strong, Perform Better. Human Kinetics: United Kingdom; 2011.

32. Harms CA, Wetter T, St Croix CM, Pegelow DF, Dempsey JA. Effects of respiratory muscle work on exercise performance. J Appl Physiol. 2000;89(1):131–8.

33. Harms CA, Wetter TJ, McClaran SR, Pegelow DF, Nickele GA, Nelson WB, et al. Effects of respiratory muscle work on cardiac output and its distribution during maximal exercise. J Appl Physiol. 1998;85:609–18.

34. Harms CA, Babcock MA, McClaran SR, Pegelow DF, Nickele GA, Nelson WB, et al. Respiratory muscle work compromises leg blood flow during maximal exercise. J Appl Physiol. 1997;82(5):1573–83.

35. Art T, Lekeux P. Work of breathing in exercising ponies. Res Vet Sci. 1989;46(1):49.33.

36. Lekeux P, Art T. The respiratory system: anatomy, physiology and adaptations exercise and training. The Equine Athlete. Rose RJ, Hodgson DR. Philadelphia: Saunders; 1994. pp. 79–128.

37. Bayly WM, Schott HC, Slocombe RF. Ventilatory responses of horses to prolonged submaximal exercise. Equine Vet J Suppl. 1995;18(S18):23–8.

38. Katz LM, Bayly WM, Hines MT, Sides RH. Ventilatory responses of ponies and horses to exercise. Equine Compar Exerc Physiol. 2005;2(4):229–40.

39. Manohar M. Blood flow to the respiratory and limb muscles and to abdominal organs during maximal exertion in ponies. J Physiol. 1986;377:25–35.

40. Bayly WM, Hodgson DR, Schulz DA, Dempsey JA, Gollnick PD. Exercise-induced hypercapnia in the horse. J Appl Physiol. 1989;67(5):1958–66.

41. Manohar M. Vasodilatory reserve in respiratory muscles during maximal exertion in ponies. J Appl Physiol. 1986;60:1571–7.

42. Manohar M. Costal vs. crural diaphragmatic blood flow during submaximal and near-maximal exercise in ponies. J Appl Physiol. 1988;65:1514–19.

43. Bayly WM, Redman MJ, Sides RH. Effect of breathing frequency and airflow on pulmonary function in high‐intensity equine exercise. Equine Vet J. 1999;31:19–23.

44. Romer LM, McConnell AK, Jones DA. Inspiratory muscle fatigue in trained cyclists: effects of inspiratory muscle training. Med Sci Sports Exerc. 2002;34(5):785–92.

45. Verges S, Lenherr O, Haner AC, Schulz C, Spengler CM. Increased fatigue resistance of respiratory muscles during exercise after respiratory muscle endurance training. Am J Physiol Regul Integr Comp Physiol. 2007;292(3):R1246–53.

46. Verges S, Renggli AS, Notter DA, Spengler CM. Effects of different respiratory muscle training regimes on fatigue-related variables during volitional hyperpnoea. Respir Physiol Neurobiol. 2009;169(3):282–90.

47. Johnson BD, Babcock MA, Suman OE, Dempsey JA. Exercise-induced diaphragmatic fatigue in healthy humans. J Physiol. 1993;460:385–405.

48. Dempsey JA, Romer L, Rodman J, Miller J, Smith C. Consequences of exercise-induced respiratory muscle work. Respir Physiol Neurobiol. 2006;151(2-3):242–50.

49. Roussos C, Macklem PT. Inspiratory muscle fatigue. Handbook of Physiology, Section 3. The respiratory system, vol 3, part 2, Mechanics of breathing. Fishman AP, Fisher AB. Bethesda, MD: American Physiological Society; 1986. pp. 511–27.

50. Guenette JA, Romer LM, Querido JS, Chua R, Eves ND, Road JD, et al. Sex differences in exercise-induced diaphragmatic fatigue in endurance-trained athletes. J Appl Physiol. 2010;109:35–46.

51. Graham R, Jammes Y, Delpierre S, Grimaud C, Roussos C. The effects of ischemia, lactic acid and hypertonic sodium chloride on phrenic afferent discharge during spontaneous diaphragmatic contraction. Neurosci Lett. 1986;67(3):257–62.

52. Jammes Y, Balzamo E. Changes in afferent and efferent phrenic activities with electrically induced diaphragmatic fatigue. J Appl Physiol. 1992;73(3):894–902.

53. Hill JM. Discharge of group IV phrenic afferent fibers increases during diaphragmatic fatigue. Brain Res. 2000;856(1-2):240–4.

54. St Croix CM, Morgan BJ, Wetter TJ, Dempsey JA. Fatiguing inspiratory muscle work causes reflex sympathetic activation in humans. J Physiol. 2000;529:493–504.

55. Rodman JR, Henderson KS, Smith CA, Dempsey JA. Cardiovascular effects of the respiratory muscle metaboreflexes in dogs: rest and exercise. J Appl Physiol. 2003;95(3):1159–69.

56. McConnell AK, Lomax M. The influence of inspiratory muscle work history and specific inspiratory muscle training upon human limb muscle fatigue. J Physiol. 2006;577:445–57.

57. Romer LM, Lovering AT, Haverkamp HC, Pegelow DF, Dempsey JA. Effect of inspiratory muscle work on periopheral fatigue of locomotor muscles in healthy humans. J Physiol. 2006;571:425–39.

58. Romer LM, McConnell AK, Jones DA. Effects of inspiratory muscle training on time-trail performance in trained cyclists. J Sports Sci. 2002;20:547–62.

59. Johnson MA, Sharpe GR, Brown PI. Inspiratory muscle raining improves cycling time-trial performance and anaerobic work capacity but not critical power. Eur J Appl Physiol. 2007;101:761–70.

60. Kraemer WJ, Fleck SJ, Evans WJ. Strength and power training: physiological mechanisms of adaptation. Exerc Sport Sci Rev. 1996;24:363–97.

61. Hershenson MB, Kikuchi Y, Tzelepis GE, McCool FD. Preferential fatigue of the rib cage muscles during inspiratory resistive loaded ventilation. J Appl Physiol. 1989;66:750–54.

62. Brown PI, Johnson MA, Sharpe GR. Determinants of inspiratory muscle strength in healthy humans. Respir Physiol Neurobiol. 2014;196:50–5.

63. Larson JL, Covey MK, Vitalo CA, Alex CG, Patel M, Kim MJ. Maximal inspiratory pressure. Learning effect and test-retest reliability in patients with chronic obstructive pulmonary disease. Chest. 1993;104:448–53.

64. Romer LM, McConnell AK. Inter-test reliability for non-invasive measures of respiratory muscle function in healthy humans. Eur J Appl Physiol. 2004;91:167–76.

65. Brown PI, Sharpe GR, Johnson MA. Inspiratory muscle training abolishes the blood lactate increase associated with volitional hyperpnoea superimposed on exercise and accelerates lactate and oxygen uptake kinetics at the onset of exercise. Eur J Appl Physiol. 2012;112:2117–29.

